# Conscious and unconscious perception of pitch shifts in auditory feedback during vocalization: Behavioral functions and event-related potential correlates

**DOI:** 10.1101/2023.10.13.562262

**Authors:** Daniel Suchý, Roozbeh Behroozmand, Henry Railo

**Affiliations:** Department of Psychology and Speech-Language Pathology, University of Turku, Finland; Turku Brain and Mind Centre, University of Turku, Finland; Speech Neuroscience Lab, Department of Speech Language and Hearing, Callier Center for Communication Disorders, School of Behavioral and Brain Sciences, The University of Texas at Dallas, 2811 N. Floyd Rd, Richardson, TX 75080, United States

**Keywords:** Speech motor control, auditory feedback, sensorimotor integration, conscious perception, neural correlates of consciousness

## Abstract

During vocalization, mismatches between expected and perceived auditory feedback are processed rapidly and automatically, suggesting that feedback control of vocalization operates unconsciously. However, whether consciousness modulates speech feedback control remains little studied. To address this question, we concurrently measured behavioral vocal responses and electroencephalography (EEG) in 30 participants while they vocalized and their auditory feedback was perturbed with individually calibrated perceptual threshold level pitch shifts. Following each vocalization trial, participants rated if they consciously detected a pitch shift in their auditory feedback. We analyzed the data on a trial-by-trial basis to test if vocal responses to pitch perturbations were modulated by conscious perception. Our results revealed that even on trials where the participants reported not noticing the pitch shift at all, a compensatory vocal response to the altered auditory feedback was generated. Conscious detection of a pitch shift was associated with an increased magnitude of vocal responses roughly 500-700 ms after the pitch shift compared to the unconscious trials. Conscious detection of the pitch shift correlated with early (Auditory awareness negativity, AAN) and late (Late positivity, LP) neural responses as indexed by the modulation of event-related potentials (ERPs). Source localization of the ERPs suggested that conscious pitch shift detection was associated with increased neural activity within the temporal, frontal and parietal cortical networks known to be involved in speech motor control. These findings emphasize the importance of investigating the role of consciousness in regulating speech feedback control, and their effect on the underlying neural and behavioral functions.

## 1. Introduction

Speech is a complex task that requires coordination of cognitive, motor, and sensory programs. Even small changes in acoustical parameters of sound can lead to meaningful changes in communication. Therefore, the process of speech production needs to be subjected to feedback monitoring that at the level of cortex involves frontal, temporal and parietal areas (Tourville & Guenther, 2011). Speech feedback control is often assumed to operate unconsciously, but the role of consciousness has rarely been probed in experimental studies. We studied if conscious perception modulates speech feedback control, and the brain mechanisms that correlate with conscious perception of pitch shifts during vocalization.

We use the term “consciousness” here to refer to the presence of subjective “phenomenal” experience; how it *feels* to hold a warm cup of coffee in one’s hand or hear one’s own voice. It is not apparent why neural activity elicited, for example, by sensory stimulation should be accompanied with any subjective experiences (Block, 1995; Nagel, 1974). In fact, research has suggested that humans can process perceptual information unconsciously (i.e., without accompanied subjective sensations about the perceived stimulus) and that unconscious perceptual information can also influence one’s behavior (Cowey, 2010; Hannula et al., 2005; Peters et al., 2017; Phillips, 2020). This leads to a series of interesting questions such as: What processes determine if a person consciously notices the perceptual information processed by her brain? What is the functional significance of conscious perception? That is, does conscious perception enable some behavioral or cognitive processes that are beyond the capacity of unconscious perceptual processes? Questions such as these are explored by a field known as consciousness science (LeDoux et al., 2020).

### 1.1. Speech feedback control

Current models of speech production (e.g., Houde & Nagarajan, 2011; Tourville & Guenther, 2011) do not distinguish between conscious and unconscious speech feedback control. Rather, the models are built around the idea that speech feedback control relies on an automatic comparison of how well the acoustic features of produced speech match the intended targets. Speech targets arise in the left-lateralized ventral premotor areas of the cortex and are executed by the motor areas. In addition to sending the feedforward motor commands, axons from the premotor cortex are assumed to relay information about the expected sensory outcome to areas such as the auditory cortex in the posterior superior temporal gyrus. This copy of the motor command is known as the efference copy (or corollary discharge). The efference copy is assumed to inhibit the neural responses to vocalizations that match the target of the motor program, and to help detect motor commands that result in mismatches between expected and produced vocalization (Behroozmand et al., 2022; Behroozmand & Larson, 2011a; Chang et al., 2013; Eliades & Wang, 2008). When a mismatch is detected, the motor areas are recruited to initiate corrections to the motor commands (Burnett et al., 1997; Hain et al., 2000).

Auditory feedback control of speech can be measured behaviorally by “perturbing” auditory feedback in real-time with artificial vocalization “errors”. In a widely used version of the paradigm, participants unexpectedly hear a pitch shifted version of their own voice during vocalization (Burnett et al., 1997; Elman, 1981; Hafke, 2008; Hain et al., 2000; Hawco et al., 2009). We call the change in vocalization that follow pitch shifts in sensory feedback the *vocal response*. This so called “frequency-altered feedback paradigm” is often used to study suprasegmental features of speech such as prosody, stress, or rhythm, and in general, the stability of pitch regulation during vocalization. These features of vocalization are crucial for spoken language, and even relatively small changes in the pitch of vocalization can lead to meaningful changes in the message conveyed (e.g., it might change the emphasis, or in tonal language it might change the meaning of the spoken word). Vocal responses to pitch perturbations are, on average, compensatory (i.e., if the perturbation increases the pitch of auditory feedback, participants typically react by lowering their voice). The vocal response is involuntary: Even when participants are asked to ignore the pitch perturbation, they still produce the vocal responses (Keough et al., 2013). Given that the onset of the vocal response is rapid (just over 100 ms), it can be assumed to be mediated by unconscious processes. However, as discussed next, previous studies that have tested the unconscious nature of vocal responding have significant methodological limitations, and consequently, the question to what extent (un)conscious perception mediates vocal responses to perturbations remains open.

### 1.2. Does consciousness modulate vocal responses?

The study by Hafke (2008) is often cited as evidence that the speech feedback control system initiates vocal responses even when the individual remains unconscious of the perturbation. Hafke (2008) showed that in trained singers, perturbations as small as 9 cents evoked vocal responses, although the participants often reported not noticing perturbations of this size. However, to show that completely unconscious perturbations produce vocal responses, the analysis should *only* include trials where participants report that they did not notice the perturbations. Hafke’s (2008) analysis also included trials where participants reported noticing the perturbation, which means that the vocal response might be contaminated by the trials where participants happen to consciously detect the pitch shifts. Another study replicated the finding of Hafke (2008) on a sample of participants without professional singing experience, but only assessed conscious perception of the perturbation by having the participants passively listen to recorded samples of perturbed or non-perturbed feedback (Xu et al., 2020). This is problematic because participants have an increased capacity to detect perturbations in their own vocalization than when the same perturbations are played back to them (Scheerer & Jones, 2018). The data by Scheerer & Jones (2018) shows that humans may be able to discriminate perturbations as small as 10– 15 cents, suggesting that 9 cent perturbations (observed to produce vocal responses in Hafke, 2008) may not completely fall short of the capacity of conscious perceptual mechanisms. Scheerer and Jones (2018) found that the differences between the thresholds for vocal responses and for conscious perception are at best minimal (5 cents) and suggested that the supposedly unconscious vocal responses to very small perturbations are instead biased by other factors such as vocal variability (i.e., participants may attribute the perturbation to the natural variability of their own voice).

Here, we aimed to test if participants initiate corrections to vocalization when they do not notice a pitch shift in their auditory feedback. If such analysis reveals that speech feedback control can be initiated solely based on unconscious perception, conscious perception could further modulate vocal responses to perturbations. In addition to the rapid vocal response component (called VR1), the vocal response may also contain a second, later component (VR2, typical onset after 300 ms). Asking participants to voluntarily control how they respond to a perturbation predominantly modifies the VR2 (Hain et al., 2000). For this reason, the VR2 is sometimes assumed to correspond to a conscious vocal response mode. The results by Franken et al. (2023) showed that when participants attributed pitch shifts to themselves, they made larger corrective responses than when they attributed the pitch shift to external factors, suggesting that conscious source monitoring of pitch shifts modulates speech feedback control. In the present study we aimed to test the behavioral function of conscious perception on vocal responses by examining if consciously noticed perturbations produce different vocal responses than perturbations which participants report not noticing. To isolate the effect of conscious perception, one needs to compare trials that differ only with respect to conscious perception while keeping physical stimulation (i.e., the pitch perturbation) constant for each participant; otherwise, the difference between conscious vs. unconscious conditions could be solely due to physical differences between the conditions.

### 1.3. EEG correlates of conscious perception of pitch shifts in vocalization

In addition to testing if conscious perception modulates speech feedback control of pitch perturbations, the second aim of the present study was to elucidate the neural processes that underlie conscious perception of pitch shifts in auditory feedback by examining event-related potentials (ERPs). Non-perturbed auditory feedback of self-produced vocalization triggers an inhibited auditory-evoked ERP compared to the same stimuli when heard through playback. This sensory suppression of self-produced speech is strongest around 100–200 ms after stimulus onset (Curio et al., 2000; Houde et al., 2002; Railo et al., 2022). When the auditory feedback the subject hears is perturbed, the amplitude of the ERPs increases, suggesting that the brain detects that the produced vocalization did not correspond to the expectation, and initiates corrections to vocalization (Behroozmand & Larson, 2011b; Heinks-Maldonado et al., 2005; Scheerer & Jones, 2014). This increase in ERP amplitude affects the ERP early (P1 wave, around 50–100 ms after stimulus onset), likely reflecting an initial unconscious detection of the vocalization pitch shift in the auditory cortex. Later processing phases (corresponding to N1–P2 waves around 100–300 ms) are assumed to correspond to more nuanced detection of the pitch shifts, and the control of subsequent corrections to vocalization.

While previous research has studied the ERP correlates of speech feedback control, the studies have not isolated the correlates of conscious perception, but rather examined how, e.g. the size of the perturbation modulates ERPs. Here, while keeping physical stimulus constant, we contrast conscious vs unconscious perception with the aim of describing what neural processes underlie individuals’ conscious perception of pitch shifts in their own voice. This type of contrastive approach is widely used in the search for “the neural correlates of conscious perception” which aims to find the neurobiological processes that enable subjective consciousness to emerge from brain activity (Crick & Koch, 1990; Dehaene, 2014). Given that physical stimulus features are kept constant for each participant, any differences between consciously noticed versus not-noticed conditions correlate with subjective conscious perception of the stimulus (see e.g., Aru et al., 2012 for more details).

Previous research suggests that conscious perception of external sensory stimuli correlates with activity in sensory, frontal and parietal cortex (Dembski et al., 2021; Koch et al., 2016; Mashour et al., 2020), although the role of the “extrasensory” areas (especially the frontal cortex) remains debated (Boly et al., 2017; Odegaard et al., 2017). ERP studies show that conscious perception of external auditory stimuli correlates with activity in two time-windows. An earlier correlate known as the Auditory Awareness Negativity (AAN) has a negative polarity and takes place around 100–250 ms after stimulus onset (Dembski et al., 2021; Eklund et al., 2021; Eklund & Wiens, 2019; Filimonov et al., 2022). This neural correlate is assumed to reflect the subjective experience of the sensory stimulus. A later correlate known as the Late Positivity (LP)—which coincides with the P3b event-related potential (ERP) wave—is often argued to reflect higher-order conscious perception (Derda et al., 2019; Jimenez et al., 2021) or post-sensory processing (e.g., decision making or behaviorally responding to a stimulus; Dembski et al., 2021; Schlossmacher et al., 2020). Other sensory modalities show similar correlates (Dembski et al., 2021).

### 1.4. Hypotheses of the present study

To vary conscious perception of the pitch perturbation while keeping its magnitude constant, we calibrate pitch perturbations for individual participants using a psychophysical staircase procedure. Whereas prior studies have often employed pitch shifts with brief duration (e.g., 200 ms; e.g., Scheerer & Jones, 2018; Behroozmand et al., 2022), following the procedure used by Hafke (2008), we here used a longer duration perturbation that lasted until the participants stopped their ongoing vocalization. We reasoned that if consciousness plays a role in speech feedback control, these effects might be best observed in situations where the pitch shift persists throughout the duration of vocalizations (i.e., when the participant notices the pitch shift, it is still present in the auditory feedback, making the participant more likely to modify their vocalization based on their conscious perception).

Following standard practice in consciousness studies, we measured conscious perception using a four-step scale commonly known as the Perceptual Awareness Scale (Overgaard & Sandberg, 2021; Ramsøy & Overgaard, 2004a; Sandberg et al., 2010). Using the graded rating scale is important because previous studies suggest that the use of binary ratings scales may contaminate the “unconscious” category with stimuli the participants faintly consciously perceived, leading to confounded results regarding unconscious processing (Koivisto et al., 2021; Overgaard et al., 2008; Phillips, 2021).

To test three preregistered hypotheses in the subsequent EEG experiment, we contrast trials where participants reported conscious perception of the perturbation with trials where they failed to detect the perturbation. First, based on earlier research (Hafke, 2008; Xu et al., 2020), we hypothesize that vocal responses are initiated even when participants report not noticing the perturbation. In addition to this unconscious mode of feedback control, we hypothesize that speech feedback control can also operate at a conscious level (e.g., Hain et al., 2000), enabling additional corrections to speech. Second, based on studies examining the neural correlates of conscious perception of external sounds (Dembski et al., 2021; Eklund et al., 2021; Eklund & Wiens, 2019; Filimonov et al., 2022), we expected that conscious perception of the perturbation correlates with an early negative amplitude difference in central electrode locations (i.e., AAN). Finally, if the ERPs in part reflect the processes that initiate vocal responses to the pitch shifts (e.g., Behroozmand et al., 2022), we hypothesized that the amplitude between ERPs and vocal responses might show a correlation.

## 2. Methods

### 2.1. Pre-registration and data availability

The present study was preregistered prior to collecting the data and running the analyses (https://osf.io/am3rq/). All procedures described in this paper conform to the pre-registration, unless otherwise specified. All computer code used to run the experiment and analyze the results are publicly available at osf.io.

### 2.2. Participants

35 participants (Mean age = 26, range: 20-36; 27 female) participated the study. All participants gave written informed consent. Sample size was determined based on previous studies, as described in the preregistration document. The participants were recruited using the university’s participant recruitment website, by advertising among psychology, speech-language pathology and biomedical students, and by word-of-mouth. The participants completed the experiment in exchange for course credit or small financial reward. The exclusion criteria for participating in the study included left-handedness, and self-reported presence of neurological, hearing, or voice disorders that would interfere with completing the experiment. A screening test of participants’ hearing was not conducted due to time constraints, and because the stimuli were individually calibrated.

We excluded five participants from the analyses. Three of them did not complete the tasks as intended. One participant reported detecting the perturbation on every trial, one varied the pitch of their vocalization systematically over the course of every trial and one reported that they were unaware of any of the large-perturbation control trials. Another two participants were excluded due to insufficient EEG quality. The research was approved by the ethics board of the Wellbeing services of the Southwest Finland and was carried out in accordance with the Declaration of Helsinki.

### 2.3. Procedure

The experimental procedure was completed in a single session, lasting up to two hours and 30 minutes. The perturbations were created using the Audapter package (Cai et al., 2008) using the phase vocoder option, and the experiment was run using the Psychtoolbox package (Brainard, 1997), both running in Matlab (version 2016B). The phase vocoder pitch-shift function shifts not just the fundamental frequency but affects the whole spectrum (Figure 1). This is perceived as a change in pitch, with no noticeable change in vowel quality. Participants’ vocalization was recorded using a Motu MicroBook IIc audio interface and a microphone (Audio-technica AT2035) positioned at about 2 centimeters distance from the mouth. The participants heard their own voice through sound-isolating in-ear earphones (Neuroscan, 10Ω). During each trial, the auditory feedback was mixed with pink noise in order to reduce the participant’s ability to hear their own voice due to insufficient earphone isolation, and bone conduction. The sound pressure level (SPL) of the auditory feedback (with the pink noise in the background) was adjusted to a level that subjectively corresponded to the intensity at which one hears their own voice without earphones. This SPL was used for all participants, except for a single participant, who considered the feedback to be too loud. We acknowledge that the lack of objective measurements about the SPL represents a limitation of the present study. That said, we feel confident that the SPL was “natural”, and that the earphones and background noise worked well in blocking the participant’s own voice “leaking” through the headphones.

**Figure 1:**
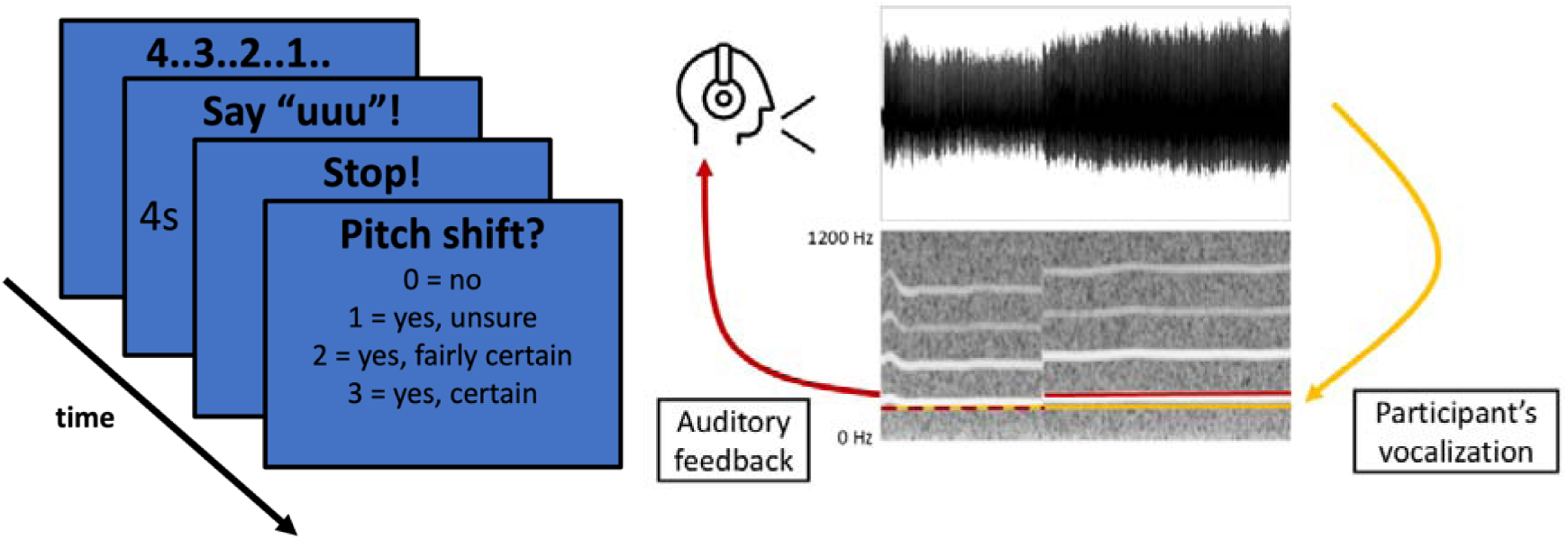
Graphical illustration of the experimental task. The participants heard their online auditory feedback through earphones while it was sometimes perturbed 1.5 seconds after the start of vocalization (upward pitch shift). In the panel on right, the yellow line denotes the true pitch of vocalization, and the red line shows the auditory feedback, which is perturbed (pitch-shifted) after 1.5 seconds. After each vocalization, the participant reported if they detected the perturbation.

Before the experiment, we instructed the participant to produce the vowel /*u*:/ of sufficient length and loudness. The participant was not instructed about the pitch of the vocalization. We did not instruct the participant on how to respond to the perturbation, other than to keep vocalizing until the preset time limit. The vowel choice was inspired by Hafke (2008). We assume that vowel choice here does not influence the results (in follow-up studies we have obtained similar results as reported here with the vowel /*a*:/). The graphical representation “uuuu” produced consistent vocalization in our international sample of participants. We chose to produce pitch perturbations rather than formant perturbations because pitch is the most frequently manipulated vocalization property in the perturbation paradigm studies (Hafke, 2008; Hain et al., 2000; Villacorta et al., 2007), and researchers typically assume the feedback control of pitch is based on unconscious processing.

At the beginning of the experiment, we first presented the participant with an interactive warm-up task, with the goal of exposing them to both unperturbed and highly perturbed feedback (+200 cents, present until the end of vocalization). During the warmup, the participants also practiced how to respond to the consciousness question that followed each trial. The warm-up continued until the desired quality of vocalization had been achieved and the participant was able to consistently identify the perturbed feedback.

After the warm-up, we presented the participant with a staircase calibration procedure, with a goal of estimating their perturbation perception threshold. The perturbation perception threshold represents a magnitude of pitch change (e.g., +20 cents) that the participant can consciously detect in a predefined proportion of trials (e.g., in 65 percent of the total trial count). We used the QUEST staircase algorithm (Watson & Pelli, 1983). The participant produced the /u/ phoneme for approximately 4 seconds. During the first 1.5 seconds of every trial, the participant was exposed to the unperturbed earphone feedback. Auditory feedback was perturbed 1.5 seconds after the beginning of vocalization, and the perturbation continued until the end of the trial. After each trial, the participant was asked to report on a 4-step scale whether they heard the change in the pitch (see below for details). The QUEST algorithm then placed the next perturbation at the value most likely to be near the specified threshold. A positive response (ratings 1–3, indicating a consciously detected perturbation) lead to a decreased perturbation magnitude being used in the next trial of the experiment, while a negative response (lowest rating) lead to an increase of perturbation magnitude. Finally, we calculated the mean of the probability density function (as recommended by King-Smith et al., 1994) and used this value as the individual perturbation perception threshold in the main phase of the experiment.

The calibration procedure had a fixed length of 50 trials, but it was re-run in case the results suggested an unreliable threshold estimate (e.g., unexpectedly high or low threshold value). All perturbations presented during the calibration as well as during the other phases of the experiment were upward, and the perturbation lasted until the end of vocalization. We initially estimated 75% perturbation detection thresholds, as it was considered an adequate trade-off between the amount of acquired unaware trials while maximizing perturbation size (to obtain clear ERPs). However, the 75% threshold resulted in a higher-than-expected amount of consciously perceived trials in the main experiment for the first participants, and therefore we lowered the threshold for later participants. In the participants included in the statistical analysis, the estimated thresholds were as follows: 75% threshold in 2 participants, 50% threshold in one participant, and 60% threshold in the remaining 27 participants. While this change is at odds with the pre-registration, it enabled us to maximize the number of aware and unaware trials per participant.

The main experiment, summarized in Figure 1, took place after the calibration session. We recorded the EEG signal only during this phase. This experiment contained 300 trials per participant, divided into blocks of 50 trials. Each block was followed by a short break. The participants were presented with perturbations at the individually calibrated intensity. Control trials without perturbations, and control trials with large perturbations (+200 cents) were presented as well. Trials without perturbations were included to verify that a participant typically reports not noticing a perturbation when it was not presented. Similarly, trials with large perturbations were used to exclude participants who systematically labelled them as ‘unaware’, and to make the task more enjoyable for the participants (by presenting, every now and then, trials where the perturbation was easily detectable). The control trials amounted to ∼33% of all trials and they were equally divided into trials with no perturbations and trials with strong perturbations. These trials were randomly distributed across the whole session (i.e., different blocks did not necessarily contain the same distribution of different trial types). The participant was instructed that they will be able to hear their unmodified voice at the beginning of each trial, and that this voice may then be modified as they continue vocalizing. After each trial, the participant was asked if they heard that the pitch of their vocalization was modified. The participant provided a response on a scale of 0 to 3, with higher numbers indicating clearer conscious perception of the perturbation. The participants were told that the question concerns the auditory signal they hear from the earphone (e.g., not how they thought they vocalized), and that the numbers on the scale corresponded to these statements: 0) No, I did not hear any change in pitch; 1) I might have heard a change in pitch, but I am unsure: 2) I am fairly certain that I heard a change; and 3) I am very confident that I heard a change. The difference between the options (0) and (1), in particular, was stressed to the participants in order to encourage a conservative criterion for the lowest alternative. In the statistical analysis the lowest alternative corresponds to the “unconscious” condition

### 2.4. Voice data processing and analysis

Participants’ vocalization was processed in Matlab version R2022b with the perturbation onset data obtained from the voice-modulation software Audapter (Cai et al., 2008). During the preprocessing of vocalization data, we first lowpass filtered the data at 500 Hz and extracted the fundamental frequency (F0) from each vocalization using Matlab’s *pitch* function. We removed the last 300 ms from the end of each trial as these contained unstable F0s. We removed trials that did not contain any vocalization, or contained vocalization that was too short, or too quiet to activate Audapter’s perturbation algorithm. Next, we determined the start time of the perturbation relative to the start of each trial, and excluded trials which did not contain at least 800 ms of perturbed phonation. We used the 200 ms before the perturbation onset as a baseline period. For every sample from 200 ms before perturbation onset until 800 ms after perturbation onset, we calculated the relative deviation of pitch in this sample from the pitch during the baseline period, using the standard equation used in auditory perturbation research (Hafke, 2008; Hain et al., 2000):

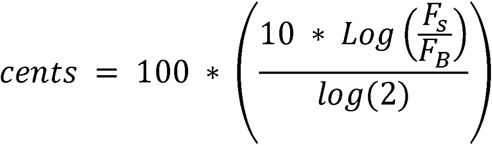

where F_s_ is the fundamental frequency of a specific audio time sample, and F_B_ is the mean fundamental frequency during the 200 ms pre-perturbation period. This calculation takes absolute units of frequency (Hertz) and translates them to relative units (cents, 100 cents = 1 semitone), which enables comparing participants and trials with varying pitch. Lastly, we excluded trials which contained vocal responses larger than 200 cents at any time point during the trial.

### 2.5. EEG data acquisition and analysis

The EEG signal was recorded from 32 scalp electrodes with NeurOne Tesla amplifier (Bittium, 2022), software version 1.5.0. Two additional electrodes were used during data collection. One was placed on the forehead (in-between the FP1 and FP2 electrodes) and served as the ground. The second one was placed on the nose and served as the reference during data collection. Sampling rate was 500 Hz. Markers that denoted participants’ keyboard responses (conscious perception ratings), and trial type were added to the EEG signal during recording. The onset of the pitch perturbation was marked in the EEG signal during data preprocessing based on Audapter data.

The EEG data was processed using EEGLAB, version 2022.1 (Delorme & Makeig, 2004) running in Matlab version R2022b. During preprocessing, channels containing a large proportion of noise and artifacts were rejected using the function *pop_rejchan*, using kurtosis and probability measures. The mean number of removed channels per participant was 6.83 (sd = 2.48). The signal was high pass filtered at 1 Hz using EEGLAB’s function *pop_eegfiltnew*. Line noise around 50 Hz was reduced using the Zapline function (de Cheveigné, 2020). Low pass filtering was not used. The data was re-referenced to average reference of all scalp electrodes. The data was epoched from 500 ms before the perturbation onset up to 1000 ms after the perturbation onset. Each trial was baseline corrected to the 500 ms time window before the perturbation onset. Next, we ran the Independent Component analysis (ICA), using the *pop_runica* function in ‘extended’ mode. After ICA, removed or missing channels were interpolated using function *pop_interp* with default parameters. Independent components were automatically labelled using ICLabel, and only those that were classified as being brain based with more than 70 percent probability were kept. Finally, we removed artifactual trials using the *pop_jointprob* function (local and global thresholds = 3). We excluded 903 trials during preprocessing (9.23 percent of the whole sample), leaving a total of 7432 trials left for the analysis described below.

### 2.6. Statistical analysis

#### 2.6.1. Vocal responses

We analyzed the vocal responses using a mass univariate analysis, that is, the effect of consciousness on the vocal response is examined at every time point before and after the onset of the perturbation. While our preregistration did not specify a mass univariate analysis, its use here is justified because it allows comparisons at high temporal resolution while controlling family-wise statistical error rate. Second, the method is suitable to situations where prior information about the timing of effects is limited, as is the case here.

While vocal responses to pitch shifts are typically in the opposite direction compared to the pitch shift, in some trials or participants, the vocal response “follows” the direction of the pitch shift. In some previous studies, the “following” responses have been excluded from the data or the two types of responses have been analyzed separately. This choice is typically motivated by the assumption that the “following” and “opposing” responses reflect two different underlying mechanisms. Following Miller et al. (2023), who showed that the vocal response distributions follow a normal distribution—suggesting that they are caused by one mechanism that displays random variation—we decided to keep both “following” and “opposing” responses in the data. This decision was further motivated by the fact that had we removed the “following” responses (or analyzed the two types of responses separately), the data distribution would have become strongly skewed, violating the normality assumption of the linear regression models used to analyze the data.

To run the mass univariate analysis of vocal responses, we have adopted a methodology typically used for mass univariate analysis of ERPs (Groppe et al., 2011; Pernet et al., 2015). First, we selected all critical trials, and ran a linear mixed effects regression model for each time point of the vocalization from 200 ms before the perturbation onset up to 800 ms after the perturbation onset. In the model, we treated the magnitude of the vocal response as the dependent variable, consciousness as a fixed-effect predictor, and the by-participant intercept as a random effect (pitch ∼ consciousness + (1|participant)). This analysis resulted in a large number of statistical tests. To account for multiple comparisons, we next ran the same series of tests with 1000 random permutations of the original trials. To do this, we randomly assigned trials to either in the aware or the unaware conditions and repeatedly ran the previously described statistical model. To identify time clusters with a significant effect of consciousness, we ran threshold-free cluster enhancement (TFCE)(Smith & Nichols, 2009) for both real regression results and the results based on the random permutations using the limo_tfce function from the LIMO EEG toolbox (Pernet et al., 2011). We selected the maximum TFCE-value (out of regression models for each time sample between −200–800 ms) for each of the random permutations and saved these to form a null-distribution to which the TFCE values of the real (i.e., not randomized) data was compared to. TFCE-values of real data higher than 97.5 of randomly permuted TFCE-values were considered significant effect of consciousness (this is equal to two-tailed alpha level of 0.05).

#### 2.6.2. ERPs

For the analysis of ERPs, we also used linear mixed-effects models on single-trial data. Unlike in the vocal response analysis, in the ERPs we expected to see two previously reported (Eklund et al., 2019, 2021; Eklund & Wiens, 2019; Filimonov et al., 2022; Schlossmacher et al., 2021) correlates: AAN (100–200 ms after stimulus onset), and LP (300–500 ms after stimulus onset). We analyzed these using two separate models, although only the analysis in the AAN time window was preregistered. In these models, we treated mean voltage during the time-period (measured on a single-trial basis) as the dependent variable, consciousness as the fixed effect predictor, and the by-participant intercept as the random effect predictor (voltage ∼ consciousness + (1|participant)). As specified in the preregistration, we only included the electrodes from the central electrode cluster (Cz, FC1, FC2, CP1, CP2) in this model (both AAN and LP are typically strongest in these electrodes; Eklund et al., 2019, 2021; Eklund & Wiens, 2019; Filimonov et al., 2022; Schlossmacher et al., 2021). To visualize the scalp distribution of AAN and LP in terms of t values, we examined how consciousness influenced ERPs by running the model separately for each channel. We performed this analysis without differentiating between following and opposing vocal responses.

To model which brain sources contribute to AAN and LP, we performed source estimation using sLoreta (standardized low resolution brain electromagnetic tomography), which solves the inverse problem of EEG source localization by generating a reference-free and polarity-independent single linear solution based on the linearly weighted sum of all scalp electrode potentials (Pascual-Marqui, 2002). Previous intracranial and brain imaging studies have validated the accuracy of the sLORETA (Mulert et al., 2004; Pizzagalli et al., 2004; Zumsteg et al., 2006). For each subject, sLORETA images of activation on cortical gray matter were generated using Montreal Neurological Institute’s standardized template (MNI152). The resolution of the present reconstruction is limited by the small number of EEG electrodes, and the lack of individual MRIs (Song et al., 2015). Statistical analysis of source activations was performed separately for early (AAN; 100–200 ms) and late (LP; 300–500 ms) time windows of ERPs by comparing the source density distributions in the conscious and unconscious conditions voxel-by-voxel using t tests. The results were corrected for multiple comparisons using permutation testing. Unlike the analysis on ERPs (which was performed on single-trial data), sLoreta analysis was based on average ERPs. This analysis was not preregistered.

## 3. Results

### 3.1. Behavioral results

The participants greatly differed in their ability to consciously detect the perturbation (Figure 2). The mean perturbation magnitude determined by the staircase procedure was 23 cents (min 1 cent, max 71 cents). The mean proportion of critical trials (i.e., trials with a perturbation whose size was adjusted using the staircase) reported as aware during the experiment was 70%, but performance varied considerably between participants (min 16%, max 95%). The overall result is in line with that of Hafke (2008), who showed that perturbation magnitudes of 19 cents are, on average, below participants’ 75% threshold of conscious perception.

**Figure 2:**
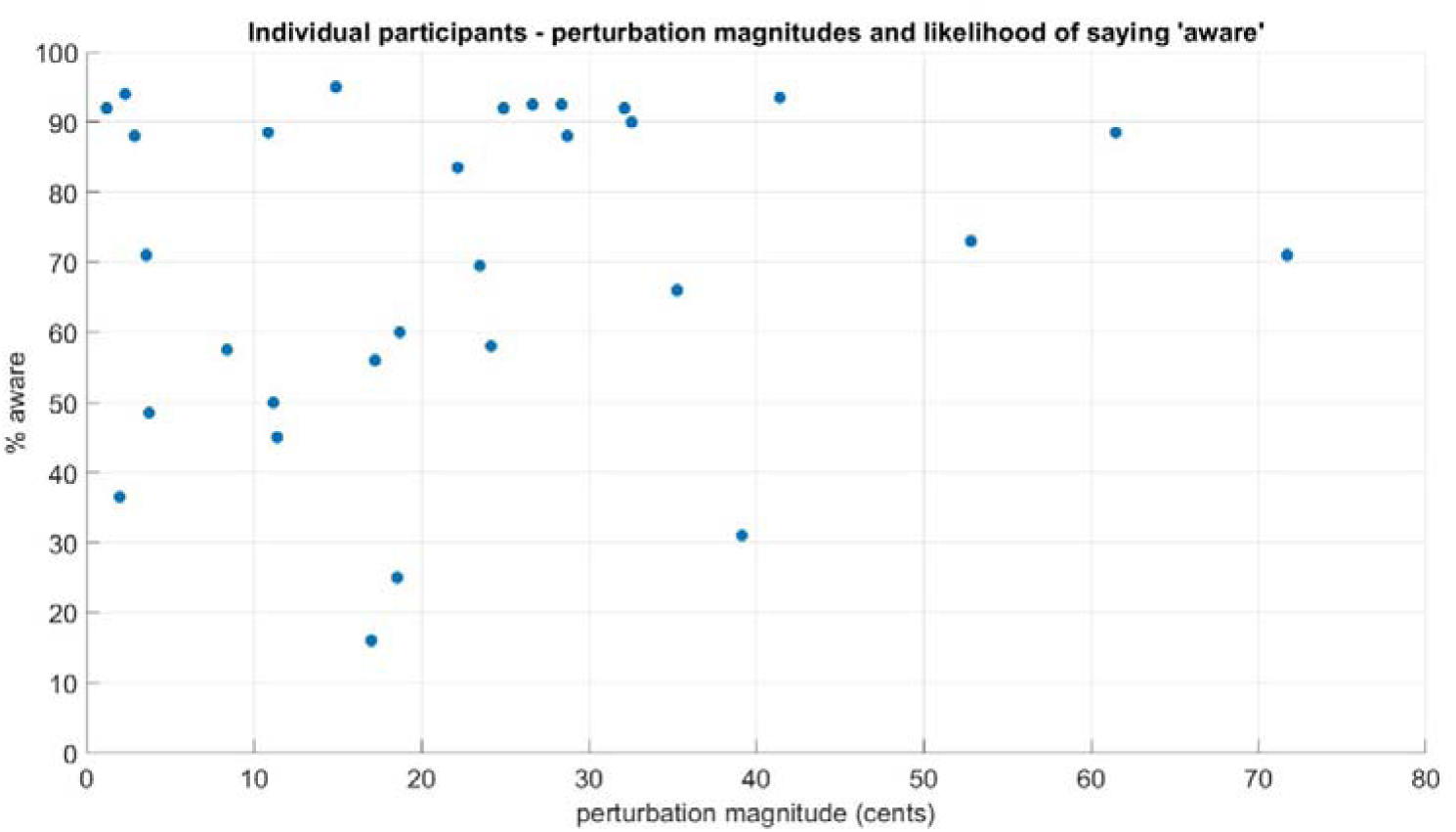
Individual participants’ thresholds for conscious perception. X axis represents the perturbation magnitude played to each participant during the experiment, obtained by the staircase method. Y axis represents the likelihood of the participant labeling a trial with an ‘aware’ rating (i.e., rating scale alternatives 1–3).

### 3.2. Vocal responses

Mean time course of vocalization during each type of trial used in the experiment is visualized in Figure 3A. To simplify the visualization of the results, conscious vs. unconscious conditions have been dichotomized for Figure 3A, although in the statistical analysis (Figure 3B) the consciousness ratings were not dichotomized. Figure 3A shows that participants also made vocal responses to threshold-level stimulus, both when they reported being conscious (ratings 1–3; yellow line) or unconscious (rating = 0; purple line) of it. Compared to the biphasic vocal response in high perturbation trials (red line; 200 cent perturbation), threshold level perturbations produced a longer duration vocal response. We quantified the statistical significance (p < .05, corrected for multiple comparisons) of the effect of conscious perception using mass univariate analysis whose result is visualized in Figure 3B. Permutation testing indicated that conscious perception of the perturbation increased vocal response magnitude in the time interval from 494 ms until 694 ms after perturbation onset (gray shaded area in Figure 3B). As shown in Figure 3C, conscious perception equally modulated both compensatory (i.e., negative) and “following” (i.e., positive) vocal responses, and altogether—in line what Miller et al. (2024) report—the responses followed a normal distribution. Vocal response magnitude increased as a function of conscious perception rating in the 494-694 ms time-window (Figure 3D) across the four rating scale steps.

**Figure 3:**
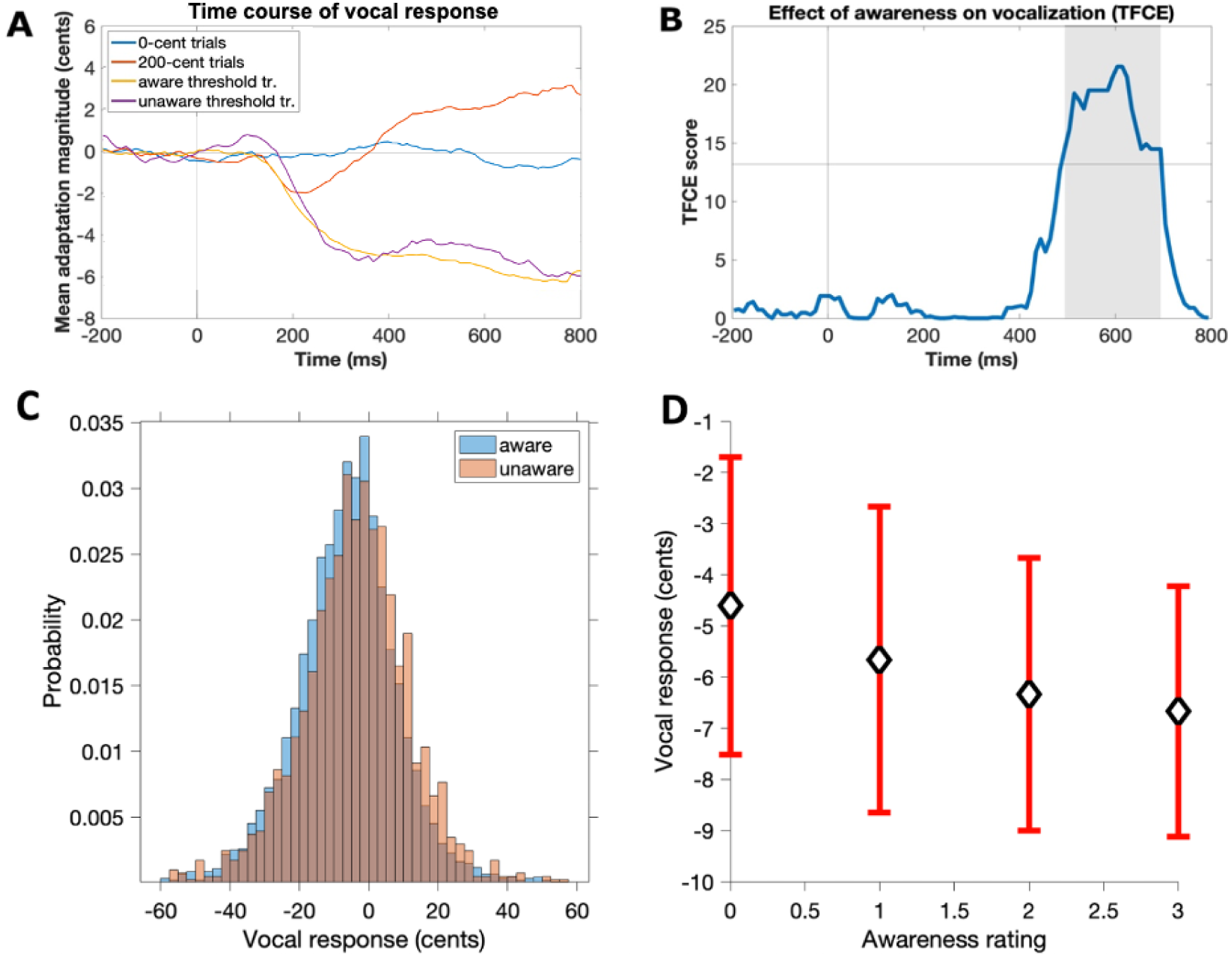
Vocal response results. (A) Mean vocal response. On the x axis, zero represents perturbation onset. On the Y axis, zero represents 200 ms pre-perturbation mean pitch. Consciousness ratings are here treated as binary for the ease of viewing (unaware = rating 0). (B) Results of the mass univariate analysis. Highlighted area represents the time interval with a statistically significant effect of consciousness, meaning that significant t-values cluster around these time points. The horizontal line represents the statistical significance threshold (p < .05, corrected for multiple comparisons). (C) Histogram of the single-trial vocal responses (pooled across different participants) 494 to 694 ms after perturbation onset separately for unaware (rating 0), and aware (ratings 1–3) trials. (D) Mean vocal response magnitude 494 to 694 ms after perturbation onset as a function of consciousness rating (pitch shift bars represent the SEM).

### 3.3. ERPs

Graphical summary of the ERP results is shown in Fig. 4. Trials rated as unconscious did not reveal a clear ERP response, while conscious trials showed early negative (100–200 ms) and late positive (300–500 ms) ERP waves (Fig. 4A). Both conscious and unconscious condition ERPs included a roughly 10 Hz oscillation (Supplementary Figure 1). We performed linear mixed effects regression tests to measure the effect of conscious perception on ERPs produced by the perturbation (Table 1). These analyses focused on two time-windows (AAN and LP). The effect of conscious perception was significant for both the early component and the late time-window, reported in Table 1 (the intercept is the amplitude in unconscious trials, and the Consciousness Estimate indicates how ERP amplitude changes as rating increases one unit). As shown in Fig. 4B and Fig. 4C, the amplitude of the early and late responses increases as a function of consciousness rating, suggesting that these correspond to AAN and LP. The scalp distributions (Fig. 4D and Fig. 4E) of the early and late time window correlates are consistent with what has been previously reported for AAN and LP, although both time windows suggest somewhat a right-lateralized activation.

**Figure 4:**
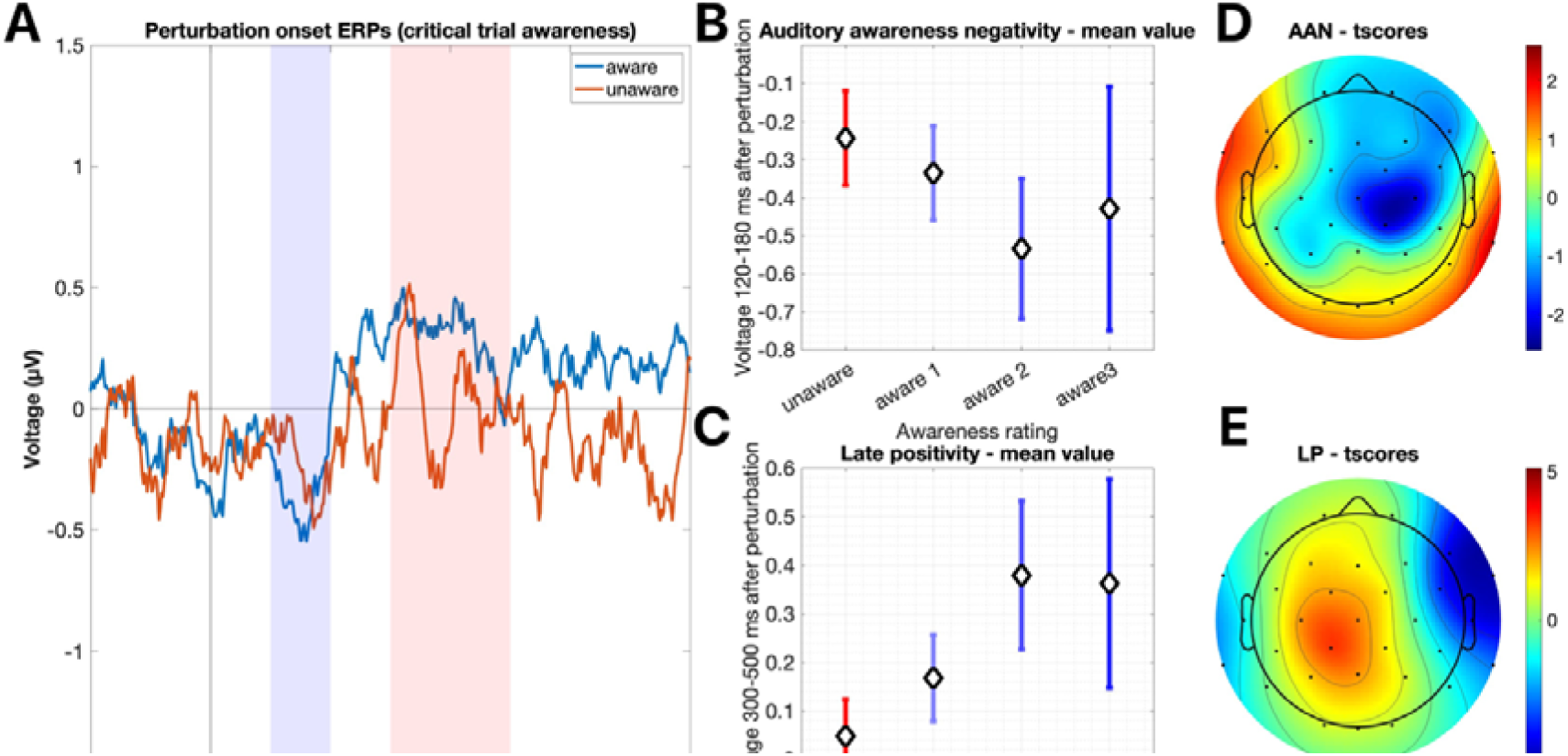
ERP results. (A) Mean ERP responses for trials rated as conscious and unconscious. Perturbation onset is at 0 ms. The consciousness rating is binarized for easier presentation (unaware = rating 0). (B) AAN amplitude as a function of consciousness. The error bars represent standard pitch shift of mean. (C) LP amplitude as a function of consciousness. The error bars represent standard pitch shift of mean. (D) Scalp distribution of AAN (map of t-values). (E) Scalp distribution of LP (map of t-values).

**Table 1:**
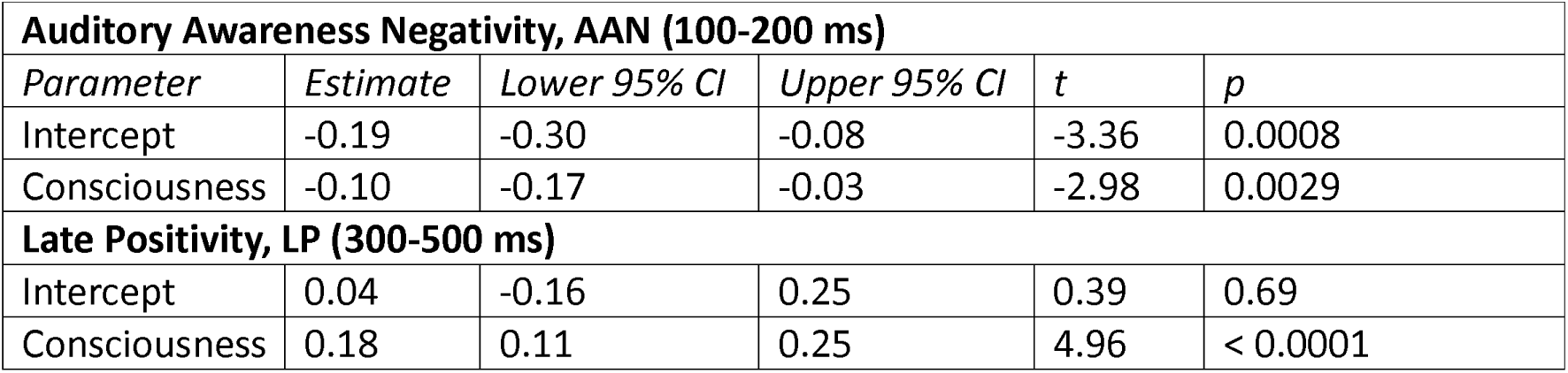
Results of linear mixed effects regression on the effects of consciousness of early and late ERPs.

### 3.4. Vocal-ERP correlation

We tested for correlations between ERP and vocal response magnitudes. We selected the early and late ERP correlates as presented above and tested if they correlate with vocal response amplitudes from (a) the same time interval, or (b) the time interval with a significant effect of consciousness (i.e., 464-694 ms), obtained from the mass univariate analysis of vocal responses (Figure 3B). We used linear mixed effects regression to compare these variables on a single-trial basis (pitch ∼ erp_amplitude + (1|participant)). None of these 4 tests showed a near significant effect of ERP magnitude on vocal response magnitude.

### 3.5. Source estimation

Figure 5 visualizes the sources that were statistically significantly modulated by conscious perception of the perturbation during AAN (top panels), and LP (bottom panel) time-windows. This analysis was performed by contrasting dichotomized consciousness ratings (i.e., rating 0 vs. ratings 1–3). Table 2 and Table 3 lists the MNI coordinates of the (statistically significant) sources maximally modulated by conscious processing of vocal feedback pitch changes in the AAN, and LP time-windows, respectively. All local maxima are reported in the tables, but if multiple significant sources exist within a single structure, only the source with the highest effect of consciousness is presented in Tables 2 and 3.

**Figure 5:**
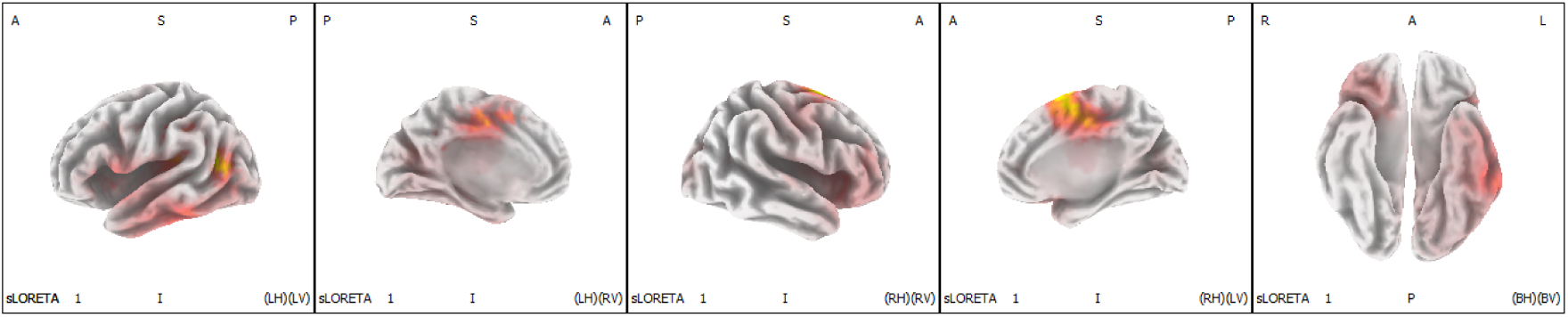
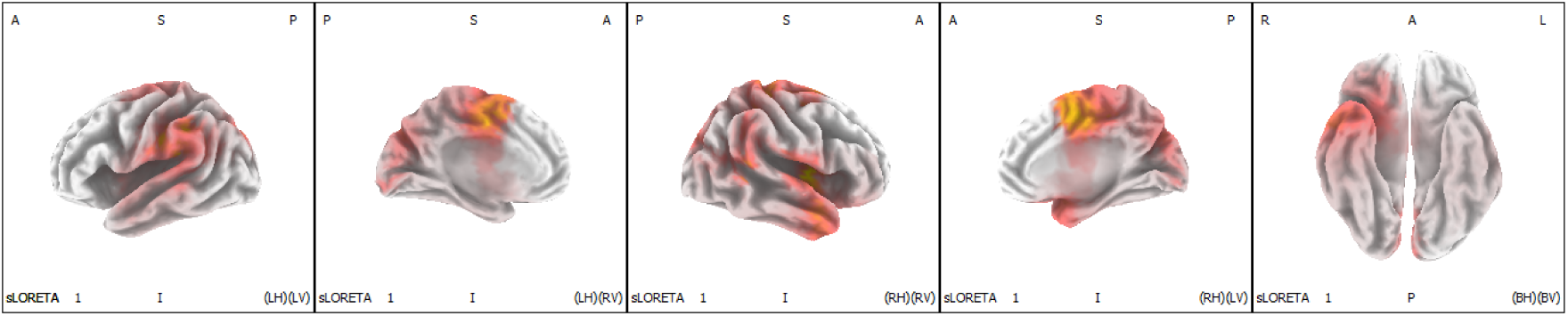
The estimated current source density maps of statistically significant (p < .05) modulation by consciousness (t test results, corrected for multiple comparisons). First row: AAN (100-200 ms post perturbation onset), second row: LP (300-500 ms post perturbation onset).

**Table 2.**
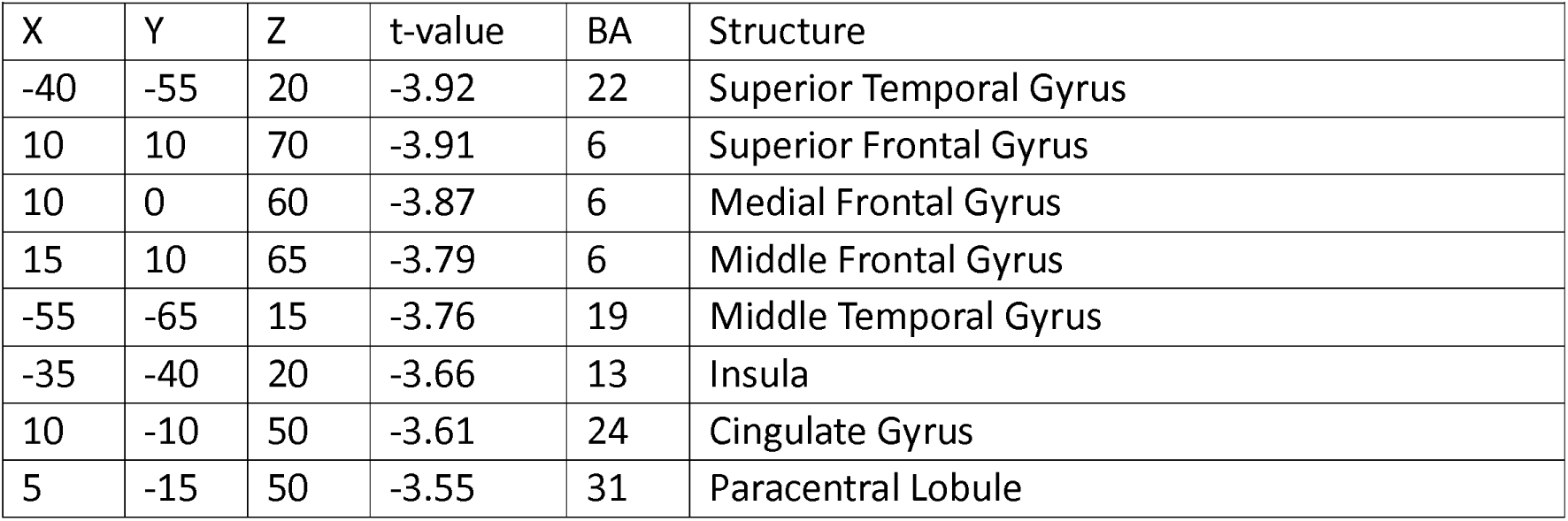
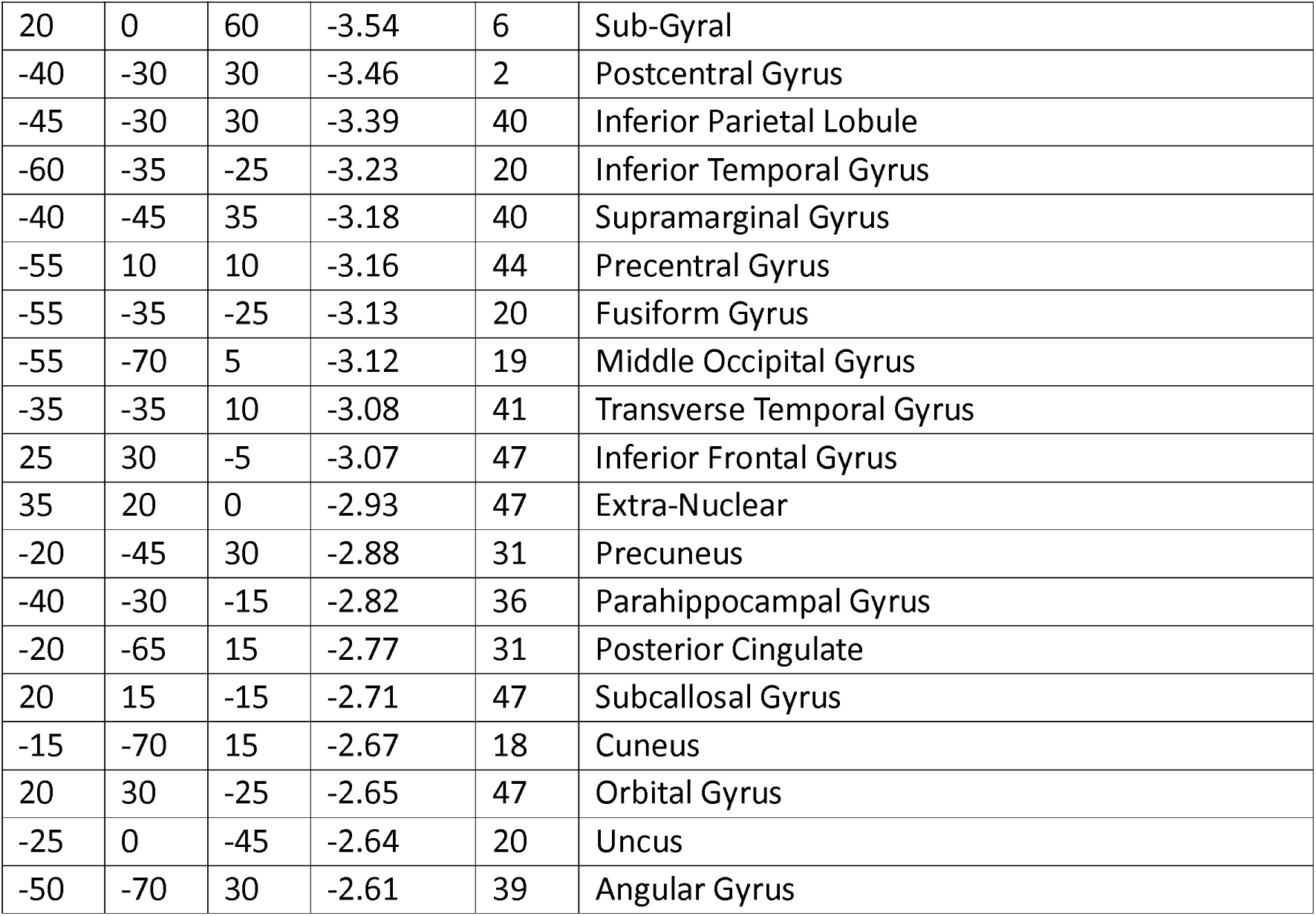
MNI coordinates of the maximum source activations in the AAN time window.

|  |  |  |  |  |  |
| --- | --- | --- | --- | --- | --- |
| X | Y | Z | t-value | BA | Structure |
| -40 | -55 | 20 | -3.92 | 22 | Superior Temporal Gyrus |
| 10 | 10 | 70 | -3.91 | 6 | Superior Frontal Gyrus |
| 10 | 0 | 60 | -3.87 | 6 | Medial Frontal Gyrus |
| 15 | 10 | 65 | -3.79 | 6 | Middle Frontal Gyrus |
| -55 | -65 | 15 | -3.76 | 19 | Middle Temporal Gyrus |
| -35 | -40 | 20 | -3.66 | 13 | Insula |
| 10 | -10 | 50 | -3.61 | 24 | Cingulate Gyrus |
| 5 | -15 | 50 | -3.55 | 31 | Paracentral Lobule |
| 20 | 0 | 60 | -3.54 | 6 | Sub-Gyral |
| -40 | -30 | 30 | -3.46 | 2 | Postcentral Gyrus |
| -45 | -30 | 30 | -3.39 | 40 | Inferior Parietal Lobule |
| -60 | -35 | -25 | -3.23 | 20 | Inferior Temporal Gyrus |
| -40 | -45 | 35 | -3.18 | 40 | Supramarginal Gyrus |
| -55 | 10 | 10 | -3.16 | 44 | Precentral Gyrus |
| -55 | -35 | -25 | -3.13 | 20 | Fusiform Gyrus |
| -55 | -70 | 5 | -3.12 | 19 | Middle Occipital Gyrus |
| -35 | -35 | 10 | -3.08 | 41 | Transverse Temporal Gyrus |
| 25 | 30 | -5 | -3.07 | 47 | Inferior Frontal Gyrus |
| 35 | 20 | 0 | -2.93 | 47 | Extra-Nuclear |
| -20 | -45 | 30 | -2.88 | 31 | Precuneus |
| -40 | -30 | -15 | -2.82 | 36 | Parahippocampal Gyrus |
| -20 | -65 | 15 | -2.77 | 31 | Posterior Cingulate |
| 20 | 15 | -15 | -2.71 | 47 | Subcallosal Gyrus |
| -15 | -70 | 15 | -2.67 | 18 | Cuneus |
| 20 | 30 | -25 | -2.65 | 47 | Orbital Gyrus |
| -25 | 0 | -45 | -2.64 | 20 | Uncus |
| -50 | -70 | 30 | -2.61 | 39 | Angular Gyrus |

**Table 3.** MNI coordinates of the maximum source activations in the LP time window.

|  |  |  |  |  |  |
| --- | --- | --- | --- | --- | --- |
| X | Y | Z | t-value | BA | Structure |
| 40 | 0 | 10 | 3.39 | 13 | Insula |
| 5 | -5 | 55 | 3.25 | 6 | Medial Frontal Gyrus |
| 5 | -5 | 50 | 3.24 | 24 | Cingulate Gyrus |
| -45 | -30 | 30 | 3.23 | 40 | Inferior Parietal Lobule |
| 5 | 5 | 60 | 3.20 | 6 | Superior Frontal Gyrus |
| 50 | -5 | 10 | 3.19 | 6 | Precentral Gyrus |
| -50 | -25 | 30 | 3.17 | 2 | Postcentral Gyrus |
| 5 | -15 | 50 | 3.08 | 31 | Paracentral Lobule |
| 65 | -45 | 15 | 3.03 | 22 | Superior Temporal Gyrus |

During the AAN time window, activity was modulated in distributed areas involving temporal, frontal, parietal, and even occipital cortex. In the temporal cortex, the effects of consciousness were strongest in the posterior part of left superior temporal gyrus and left inferior temporal cortex. Inferior frontal cortex and insula were activated bilaterally. Activations were also present in areas corresponding to motor and somatosensory areas, and strong activity was present especially in bilateral paracentral lobule and middle part of the cingulate cortex.

In the LP time window, bilateral paracentral lobule and middle cingulate cortex again showed strong modulation by conscious perception. Activity in the superior temporal gyrus was also modulated by conscious perception bilaterally. Conscious perception related activity is also present near the boundary of temporal and parietal cortex (this effect is somewhat left lateralized), in addition to more posterior parietal/occipital effects. Unlike in the AAN time window, the LP time-window showed considerably weaker effects in the frontal lobe.

As discussed further below, the results of the source localization should be interpreted cautiously. To try to validate the source reconstruction results, we ran control analyses where we contrasted the high-perturbation against no-perturbation (i.e., control) trials. While the number of trials included in these analyses are relatively low (because these control conditions were mainly included as behavioral controls), we reasoned that this contrast should reveal activation in cortical areas known to be involved in auditory feedback control of speech. The results revealed activations in frontal, temporal and parietal areas, largely overlapping with the source localization results reported above (see Supplementary Material for the results).

## 4. Discussion

Using an experimental approach that allowed us to contrast instances where participants consciously perceive pitch shifts in their auditory feedback with trials where the same pitch shift goes unnoticed, we characterized the behavioral and neural correlates of conscious perception of pitch shifts in one’s own voice. Our results showed that vocal responses to pitch perturbations were elicited even when participants failed to consciously notice the pitch shift, but conscious perception of the pitch shift in auditory feedback correlated with stronger vocal responses at late time windows (around 500–700 ms after the pitch shift onset). Our finding that consciousness modulated vocal responses suggests that models of speech feedback control need to explain how unconscious and conscious modes of speech feedback control differ in terms of their mechanisms and neural correlates. Our results indicate that conscious perception of a pitch shift in one’s own vocalization correlated with ERPs in early (AAN) and late (LP) time windows. Source reconstruction results suggest that during both these time windows, activity in distributed cortical areas correlated with conscious perception.

### 4.1. Effect of consciousness on vocal responses

The present results confirm the hypothesis that vocal responses to pitch perturbation are mediated by unconscious processes (Hafke, 2008; Keough et al., 2013; Xu et al., 2020). Because consciousness correlated with vocal responses only in relatively late time windows, our result is consistent with the proposal that the early vocal response component (VR1) is determined by unconscious processes, whereas the later component (VR2) can be modulated by consciousness (Hain et al., 2000).

The finding that consciousness only modulated late vocal responses aligns with current models of conscious perception. The early, unconscious part of vocal response is assumed to be mediated by a rapid, serial control circuit: mismatch between intended and produced vocalization activates pitch shift units in auditory cortex, which communicate the pitch shift to frontal cortex to rapidly initiate corrective movements. While this circuit compares auditory feedback to the learned, feedforward motor predictions (based on DIVA; e.g., Tourville & Guenther, 2011), the circuit itself is serial (i.e., “feedforward”), and does not involve the activation of recurrently activated cortical feedback loops. Theories of consciousness assume that this type of serial feedforward neural activity is not associated with conscious perception, although such activity can nevertheless trigger behavioral responses (Hurme et al., 2017; Lamme & Roelfsema, 2000). Consequently, consciousness likely only correlated with late vocal responses because conscious perception requires the activation of sustained, recurrent brain activity via feedback connections. This suggests that conscious perception of pitch shifts in vocalization may enable more precise adjustments to speech motor commands (enabling, e.g., more precise control of prosody). Within the DIVA model, the more precise control of vocalization based on consciously perceived pitch shifts in auditory feedback could indicate that (the feedforward) motor targets are updated based on the consciously detected pitch shift, whereas the rapid, unconscious auditory speech feedback control is enabled by “adding” information about the pitch shift to the ongoing motor commands (e.g., Guenther, 2016). As discussed in section 4.3. *The role of consciousness in speech feedback control*, we emphasize that the present findings are correlative and therefore do not demonstrate that the participants volitionally modified their vocalization based on the consciously detected pitch shifts.

In the present study, consciousness was associated with relatively minor changes in the vocal response magnitude. This is understandable, given that we did not instruct participants to modulate their vocalization. In our results, clearly consciously perceived perturbations (i.e., rating 3) were associated with almost 2 cents larger vocal responses than unconscious perturbation (i.e., rating 0; Figure 3D), corresponding to over 10% increase in vocal response magnitude relative to (average) perturbation magnitude. The effect of consciousness on vocal response magnitude is more significant when one compares it to how much vocal responses change as a function of perturbation magnitude. In the study by Scheerer & Jones (2018) increasing the perturbation from 0 to 40 cents increased the early vocal response altogether about 10 cents, but the late response only little over 3 cents. Although direct comparisons between studies is difficult due to methodological differences, this comparison suggests that the effect size of the influence of conscious perception on vocal response magnitude is meaningful. More generally, in the present study, the vocal response seems to have a somewhat longer duration when compared to previous studies. We attribute this difference to the fact that in our study, the perturbation lasted longer than in many previous studies.

Our results are in line with previous research (Hafke, 2008; Scheerer & Jones, 2018), suggesting that participants can detect 20 cent pitch perturbations, on average, with roughly 75% accuracy. However, our results also suggest that there is a lot of individual variation the perceptual thresholds, and some participants had very small perceptual thresholds. It is possible that the large individual variation we observed here is due to methodological limitations. Measuring the thresholds turned out be challenging. We assume that a two-alternative forced-choice paradigm that minimizes the influence of criterion shifts in perceptual decisions would enable more precise measurement of detection thresholds when compared to the single stimulus rating task employed in the present study (because two-alternative forced-choice methodology does not require setting an internal standard) (Morgan et al., 2012). Because the perturbations had a long duration in the present study, it could also be argued that some participants could detect the perturbation based on some confounding information such as beat tones (a vibration caused by the two simultaneous tones that differ slightly in frequency). We used sound-isolating in-ear earphones and embedded auditory feedback in background noise to eliminate the beat-tones. However, it is difficult to verify that we were able to eliminate beat tones completely. Finally, the difficulty in measuring perceptual threshold may be due the fact that with low perceptual thresholds, detection of the perturbation is significantly influenced by the natural variation of pitch during vocalization (i.e., participant detects natural shifts in the pitch of their voice but reports this as a “perturbation”).

### 4.2. Neural correlates of consciously perceiving pitch shifts in auditory feedback

The ERP observed in the conscious condition in the present study resembles previously reported ERPs to pitch perturbations (Behroozmand & Larson, 2011a; Scheerer & Jones, 2018a). The ERPs we observed lack the classical P1-N1-P2 pattern often observed in perturbation studies likely due to the small size of the perturbations. Scheerer & Jones (2018) observed that only perturbation magnitudes larger than 20 cents displayed the P1-N1-P2 pattern.

The ERP correlates of conscious perception in the present study reveal a similar pattern of correlates as conscious perception of purely “external” auditory stimuli (i.e., stimuli not-related to the individual’s own behavior), although, unlike in previous studies with “external” stimuli, the scalp distributions of AAN and LP are somewhat right lateralized (Dembski et al., 2021; Eklund et al., 2021; Eklund & Wiens, 2019; Filimonov et al., 2022). The primary sources of AAN have been in previous studies (that use external stimuli) localized to posterior superior temporal cortex (Dykstra et al., 2016; Eklund et al., 2019; Gärtner & Gutschalk, 2021; Wiegand & Gutschalk, 2012), although some results also report activity in frontal cortex (Dykstra et al., 2016; Eklund et al., 2019). Intriguingly, the present source localization results suggest that activity in distributed cortical areas contributed to AAN, in addition to sources in the left auditory cortex. Frontal activations were largely lacking in LP time window in the present study. This contrasts the typical interpretation that LP underlies “accessing” conscious contents for task-relevant processing through mechanisms that involve the frontal cortex (Pitts et al., 2012; Schlossmacher et al., 2021).

The present source localization is limited in resolution due to the small number of electrodes whose locations were not digitized, and the use of template brain anatomy (Song et al., 2015). Confounds associated with speech (e.g., activity observed near the temple regions could be contaminated by vocalization) and oscillatory confounds in ERPs (see below for discussion) could also bias the results of source reconstruction. Yet, the observed sources are encouragingly consistent with the neural basis of speech production (Behroozmand et al., 2022; Tourville et al., 2008; Toyomura et al., 2007). The control source reconstruction analyses are also consistent with the circuitry underlying speech production, although it should be noted that they are more widespread than the neural circuits assumed to underlie auditory feedback control of speech. Next, we first discuss the source activations in light of neural correlates of consciousness. After this, in the next chapter, we discuss how the observed sources activations may relate to neural mechanisms of speech motor control.

The widespread nature of the sources correlating with conscious perception of the perturbation in the AAN time window raises the possibility that conscious perception of pitch shifts in one’s own vocalization (and possibly the subsequent modifications to vocalization) is mediated by activity in frontal and parietal areas. Given that the sources in AAN time-window strongly overlapped with areas known to be involved in speech feedback control (Guenther, 2016), the present results suggest that that the same circuit that underlies speech motor control may also mediate conscious perception of the pitch shifts in vocalization. Put in terms frequently used in consciousness research, the present results suggest that participants may be able to consciously “access” information processed by the speech feedback control circuitry.

Based on theories of consciousness, conscious perception of a pitch shift in vocalization could require, for example, that information about the detected pitch shift is integrated into a singular network with its own cause-effect power (often correlating with parietal activity; Tononi et al., 2016), that sensory activation “ignites” strong, sustained activation in distributed areas (Mashour et al., 2020), that higher-order frontal mechanisms take the auditory pitch shift information as content (Brown et al., 2019), or that auditory cortex activates feedback connections to form recurrently communicating neural coalitions (Lamme, 2010; Lamme & Roelfsema, 2000). By and large, the present results are consistent with any of these theories, and future research is required to test which theories best capture the neural processes that correlates with the perception of pitch shifts in one’s own auditory feedback.

The observed correlates of consciousness could reflect processes that do not directly produce conscious perception of a pitch shift in vocalization, but rather “just correlate” with it (i.e., the correlates may reflect “prerequisites” or consequences of conscious perception; Aru et al., 2012). For example, the correlates observed in the present study likely include neural correlates of attention to one’s vocalization as the participants closely attended to their voice to detect the perturbation. Attending one’s own voice modulates ERPs (Hu et al., 2015; Liu et al., 2018) in the same time-windows as consciousness in the present study, and additional research is needed to tease apart the overlap between consciousness and attention. Research suggests that attention and conscious perception interact but can also be dissociated from each other (Koivisto et al., 2005, 2006; Pitts et al., 2018).

In the present study, detecting the perturbations required participants also to separate the artificial pitch shift from normal variation in the pitch of their vocalization. This type of conscious monitoring of the source of the pitch shift is known to modulate vocal response amplitudes (Franken et al., 2023; Subramaniam et al., 2018). The neural correlates of source monitoring of vocalization remain unknown, but based on studies in other modalities, it may be mediated by efference copy circuits similar to speech feedback control (David et al., 2008), and activate frontal and parietal regions such as the insula and temporoparietal junction (Kühn et al., 2013; Zito et al., 2020).

The ERPs observed in the present study contained an oscillation (roughly 10 Hz, i.e., alpha frequency; Fig. 4A and Supplementary Fig. 1). We hypothesize that vocalization onset might have triggered a phase-locked oscillation at roughly 10 Hz frequency. Because the perturbations were presented at a constant delay after vocalization onset in the present study, such phase-locked oscillation might then remain in the averaged ERP. If correct, this interpretation entails that such phase-locked oscillatory activity might also play a role in speech motor control (e.g., oscillatory action control network activity; Dosenbach et al., 2025; or tracking the sensory consequences of speech; Cheung et al., 2016), and consequently such oscillation would merit further investigation. Alternatively, the confound could also reflect e.g., muscular artifacts associated with vocalization. While the origin of the oscillation remains open, we acknowledge that it represents a limitation, and the results of the present study remain to be verified by later studies.

### 4.3. The role of consciousness in speech feedback control

The sources that correlated with conscious perception overlap with the circuitry of speech feedback control. Vocalization pitch shifts are assumed to be detected in higher-order auditory cortex in posterior superior temporal gyrus (pSTG; which receive efferent copies from the motor cortex)(Hickok, 2022). Consistent with this, in the present study, left pSTG was activated during AAN and LP time windows. In the DIVA model, auditory pitch shifts are assumed to be translated to motor coordinates in the right inferior frontal gyrus (rIFG; Tourville et al., 2008; Toyomura et al., 2007). Activity in this area correlated with consciousness in the AAN time-window. Activity in the left IFG also correlated with consciousness during AAN. This area is an integral part of speech production (Guenther, 2016), and it has been suggested to contribute to speech feedback control by regulating activity in sensorimotor areas (Behroozmand et al., 2022). Activity in the left IFG has also previously been reported to correlate with AAN in (Dykstra et al., 2016). Also consistent with present results, brain imaging results suggest that right insular cortex activity correlates with pitch shifts in speech feedback (Toyomura et al., 2007). Bilateral paracentral activity (i.e., supplementary motor area), and the adjacent Cingular cortex correlated strongly with consciousness in both AAN and LP time windows. Both of these areas are activated during speaking (Guenther, 2016; Picard & Strick, 1996), and their stimulation or lesions results in speech-arrest or mutism (Filevich et al., 2012; Guenther, 2016). In the LP time-window, the area near the boundary between parietal and temporal areas correlated with consciousness. This corresponds to area referred to as spt in the dual-stream model, assumed to underlie sensory-to-motor transformations (Hickok, 2022). Activation in nearby areas during LP also includes the angular gyri, which have been reported to correlate with processing altered auditory feedback (Behroozmand et al., 2022).

What is the relationship between the observed correlates of conscious perception of the perturbation and vocal control? Broadly, there are two not mutually exclusive explanations. First, the observed correlates of consciousness may have causally modulated vocal responses. Given that both AAN and LP preceded the time-window when vocalization was modulated by consciousness, this conclusion is in line with the observed data. According to the standard interpretation, conscious control of behavior is mediated by processes taking place in the LP time window, although some evidence suggests that already early correlates of consciousness could mediate behavioral changes (Railo et al., 2015). Given that source reconstruction suggested that activity in frontal and parietal areas correlated with conscious perception in the AAN time window, our findings are consistent with the interpretation that these processes are part of the neural mechanisms that initiated corrections to vocalization when the pitch shift was consciously noticed. However, neither AAN nor LP amplitude predicted vocal response magnitudes, suggesting that neither of ERP correlates of consciousness causally drive vocal responses.

The alternative explanation is that conscious perception of the perturbation (and therefore its neural correlates) did not causally influence the vocal responses. Based on this interpretation, a “third factor” explains the observed association between conscious perception and vocal responses. It could be, for example, argued that participants consciously detected the perturbations only on trials where they succeeded to selectively attend to the perturbation—and the same attentional selection possibly mediated amplified vocal responses. This interpretation is consistent with the fact that attention could modulate vocal responses (Hu et al., 2015; Liu et al., 2018).

Even if consciousness may not necessarily causally influence the feedback control of speech, it could play some other functional role in speech motor control and causally mediate adjustments to feedforward motor commands. For example, it is reasonable to assume that a person who consciously notices unwanted acoustic features in her own voice (e.g., a patient with voice deficits), might be able to learn optimal motor commands faster than a person who does not notice such features. If this is the case, consciousness might mediate changes that happen at longer time scales (e.g., across trials; Hantzsch et al., 2022) than examined in the present study. Finally, while in the present study we examined a very basic and automatic feature of speech motor control, future experiments with more complex speech tasks (e.g., spoken sentences where prosody has to be controlled deliberately to convey a precise meaning), might be better suited to uncover the functional role of consciousness in speech motor control.

### 4.4. Conclusion

Our results suggests that feedback control of speech functions in two different modes: A rapid unconscious mode, and a slower conscious mode that builds on the unconsciously determined vocal response. Activity in distributed cortical areas, including temporal, frontal and parietal cortices correlated with conscious perception of pitch shifts in vocalization, overlapping with circuits that underlie speech feedback motor control. Lack of understanding about how humans become conscious of their own voice, and how this enables them to voluntarily modulate their speech represents a significant gap in our understanding of the mechanisms of speech. Our results represent the first steps towards understanding this process, but significant questions remain open: What are the functional contributions of different cortical areas to conscious perception of pitch shifts in one’s own vocalization? Through what mechanism does conscious perception modulate vocalization? Could biases in how one consciously perceives one’s own voice lead to clinically meaningful speech deficits?

## 5. Author contributions

H.R. and D.S. designed the study. D.S. collected the data, and performed the statistical analysis, and visualizations. H.R. supervised data preprocessing and analysis, and R.B. supervised the source reconstruction and analysis. D.S. and H.R. wrote the first draft of the paper. All authors provided feedback and modifications to the manuscript and approved the final version of the manuscript.

## 6. Acknowledgements

This research was funded by the Research Council of Finland (grant number 351109).

## Supplementary Material

*Suchý et al*.

**Supplementary Figure 1.**
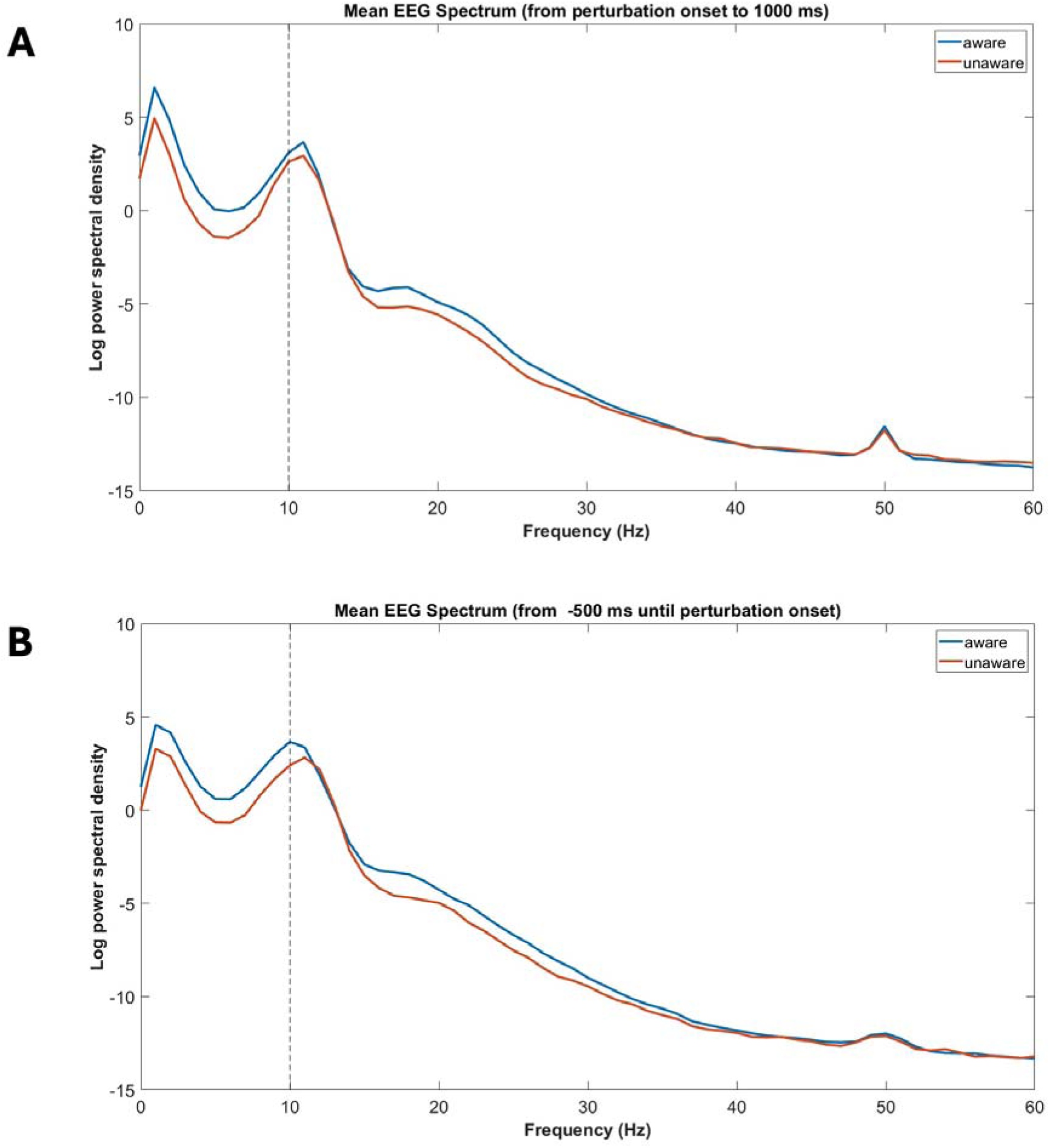
Power spectral density of the grand average ERP reported in manuscript (Fig. 4) A) after and B) before perturbation onset. The blue line depicts the aware and the red line the unaware condition.

**Supplementary Figure 2.**
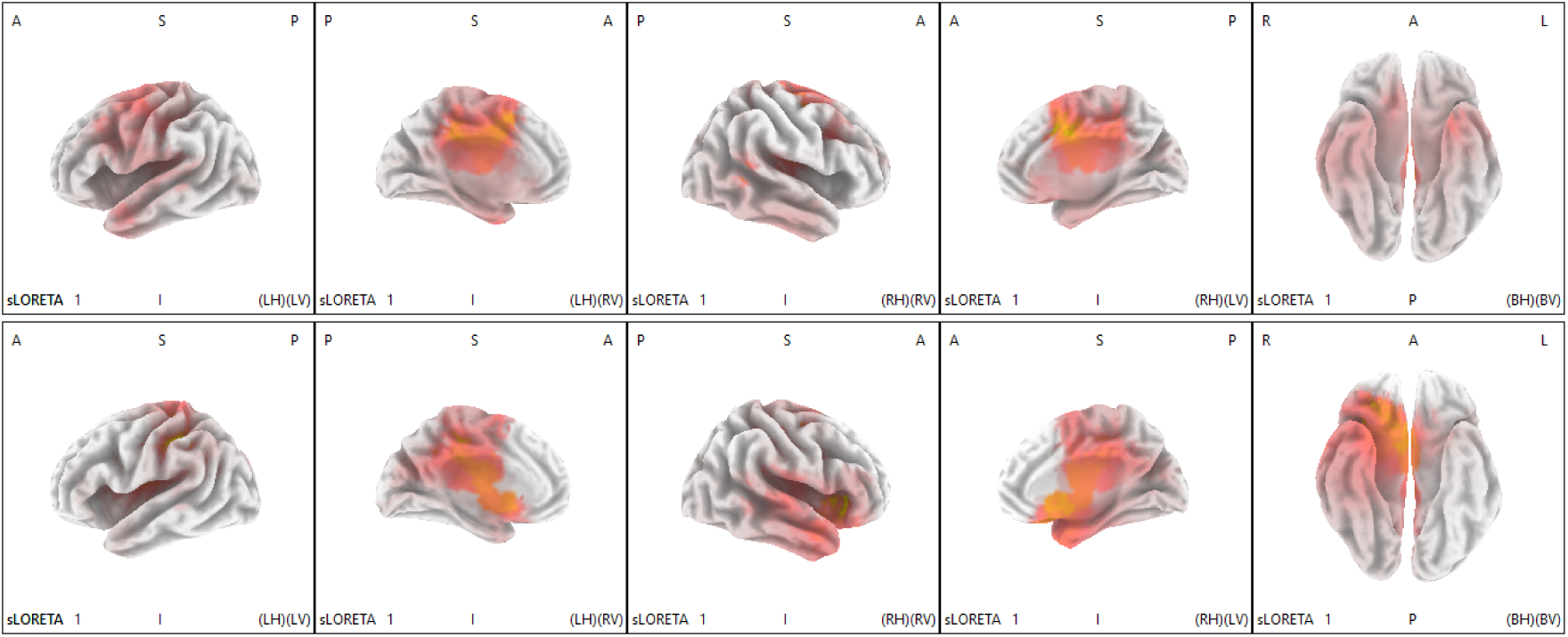
Control source localization results visualized. The heatmaps show the estimated current density maps of statistically significant (p < .05) modulation by +200 cent pitch perturbation in auditory feedback. First row: early time-window (100–200 ms post perturbation onset), second row: late time-window (300–500 ms post perturbation onset).

**Supplementary Table 1.**
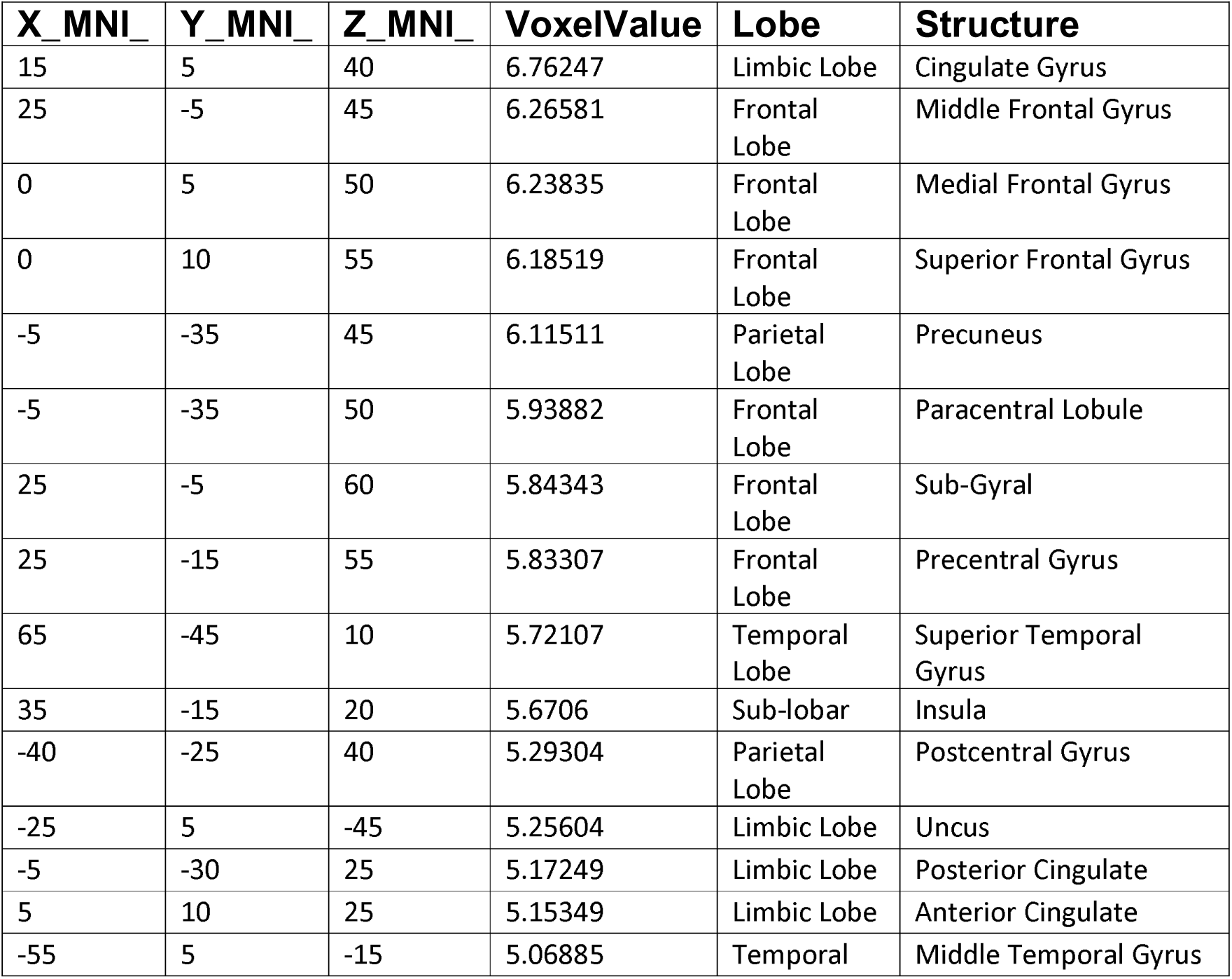
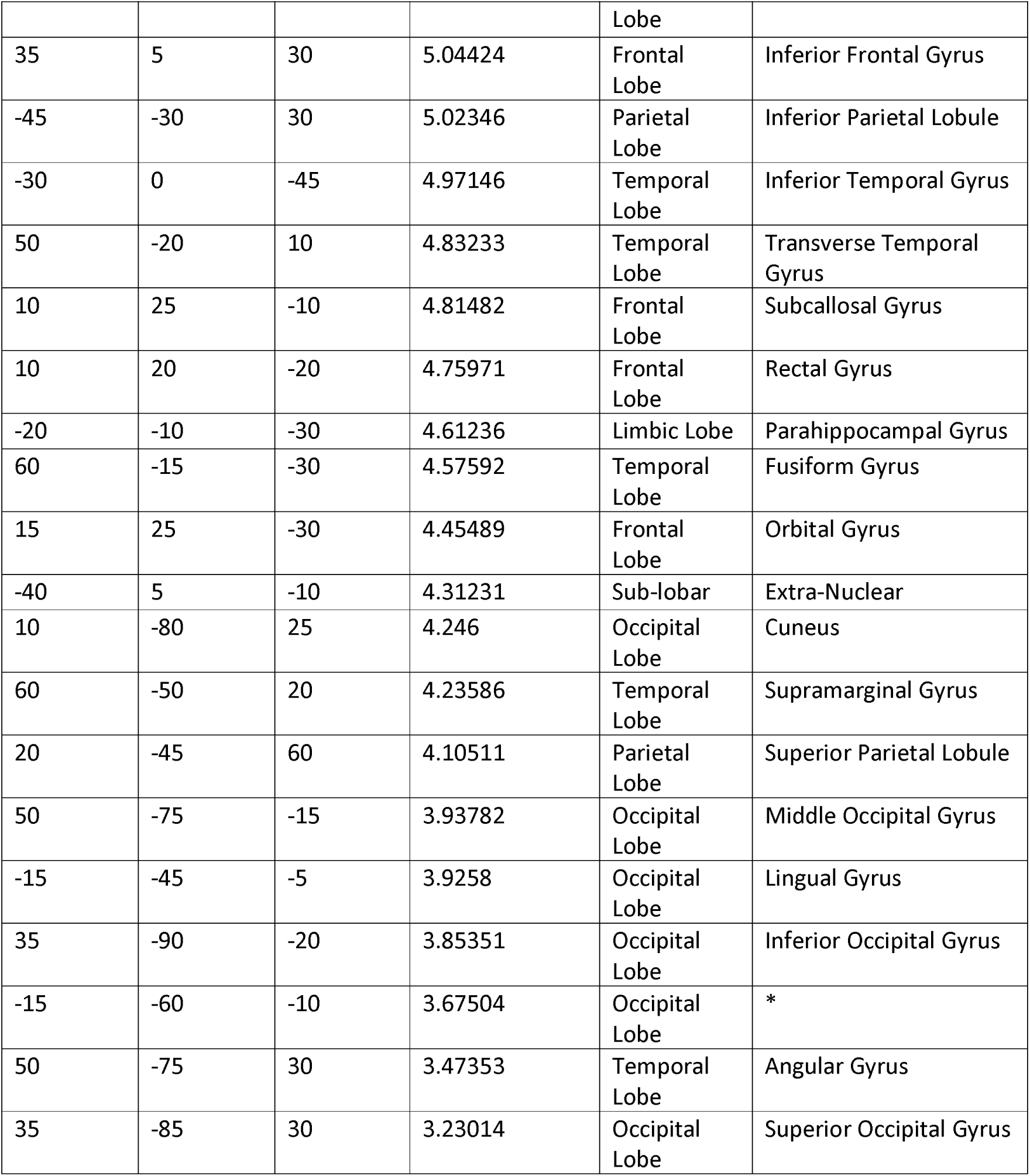
MNI coordinates of the maximum source activations in the early time-window.

**Supplementary Table 2.** MNI coordinates of the maximum source activations in the late time-window.

|  |  |  |  |  |  |
| --- | --- | --- | --- | --- | --- |
| X_MNI_ | Y_MNI_ | Z_MNI_ | VoxelValue | Lobe | Structure |
| -35 | -35 | 40 | 7.87954 | Parietal Lobe | Inferior Parietal Lobule |
| -35 | -30 | 40 | 7.82762 | Parietal Lobe | Sub-Gyral |
| 30 | 25 | 0 | 7.77795 | Sub-lobar | Insula |
| 25 | 30 | -5 | 7.76607 | Frontal Lobe | Inferior Frontal Gyrus |
| -35 | -25 | 40 | 7.55905 | Frontal Lobe | Postcentral Gyrus |
| 30 | -20 | 45 | 7.44059 | Frontal Lobe | Precentral Gyrus |
| -20 | -30 | 40 | 7.43841 | Limbic Lobe | Cingulate Gyrus |
| 35 | 20 | 0 | 7.4022 | Sub-lobar | Extra-Nuclear |
| 5 | 5 | -5 | 7.2625 | Limbic Lobe | Anterior Cingulate |
| 10 | 20 | -10 | 7.20209 | Frontal Lobe | Medial Frontal Gyrus |
| 20 | 25 | -15 | 7.11821 | Frontal Lobe | Middle Frontal Gyrus |
| 15 | 25 | -25 | 7.02934 | Frontal Lobe | Orbital Gyrus |
| 5 | 10 | -10 | 7.01546 | Frontal Lobe | Subcallosal Gyrus |
| 10 | 20 | -20 | 6.93431 | Frontal Lobe | Rectal Gyrus |
| -15 | -35 | 50 | 6.91638 | Frontal Lobe | Paracentral Lobule |
| -40 | -45 | 35 | 6.87903 | Parietal Lobe | Supramarginal Gyrus |
| -20 | -45 | 35 | 6.83787 | Parietal Lobe | Precuneus |
| 45 | 5 | -5 | 6.74605 | Temporal Lobe | Superior Temporal Gyrus |
| 15 | 0 | -15 | 6.6997 | Limbic Lobe | Parahippocampal Gyrus |
| 20 | 5 | -25 | 6.59555 | Limbic Lobe | Uncus |
| 60 | 5 | -15 | 6.54326 | Temporal Lobe | Middle Temporal Gyrus |
| -5 | -30 | 25 | 6.46609 | Limbic Lobe | Posterior Cingulate |
| -5 | 5 | 70 | 6.168 | Frontal Lobe | Superior Frontal Gyrus |
| 65 | -15 | 10 | 6.15333 | Temporal Lobe | Transverse Temporal Gyrus |
| 30 | 0 | -45 | 6.14149 | Temporal Lobe | Inferior Temporal Gyrus |
| 40 | -10 | -30 | 5.68316 | Temporal Lobe | Fusiform Gyrus |
| -10 | -70 | 30 | 5.58324 | Occipital Lobe | Cuneus |
| -20 | -45 | 60 | 5.44843 | Parietal Lobe | Superior Parietal Lobule |
| 15 | -45 | -5 | 4.74556 | Occipital Lobe | Lingual Gyrus |
| -30 | -65 | 35 | 4.38372 | Parietal Lobe | Angular Gyrus |
| 15 | -60 | -10 | 4.38348 | Occipital Lobe | * |
| 35 | -90 | -20 | 4.20848 | Occipital Lobe | Inferior Occipital Gyrus |
| 20 | -90 | -15 | 4.13788 | Occipital<br>Lobe | Middle Occipital Gyrus |
| 40 | -80 | 25 | 3.55079 | Occipital<br>Lobe | Superior Occipital Gyrus |

